# The human gut microbiome activity is resilient and stable for up to six months: a large stool metatranscriptomic study

**DOI:** 10.64898/2026.03.09.710644

**Authors:** Ryan Toma, Lan Hu, Nan Shen, Eric Patridge, Robert Wohlman, Guruduth Banavar, Momchilo Vuyisich

## Abstract

The human microbiome influences health and disease through diverse biochemical and functional outputs (e.g. enzymes, structural proteins, metabolites, and other cellular components) that affect nearly every aspect of human physiology. Metatranscriptomics (MT), an unbiased RNA sequencing approach, is a high-throughput and high-content method that quantifies both gut microbial taxonomy and active biochemical functions. Because microbial community composition and gene expression are dynamic, understanding temporal variation in the gut metatranscriptome across multiple time scales is essential. Here, we report the temporal dynamics of gut microbiome species and functions using a large cohort (n=6,157) with a clinically validated stool MT test. We quantified microbiome stability from hours to years and assessed taxonomic and functional resilience to major luminal perturbations, such as colonoscopy bowel preparation. Longitudinal analyses of samples collected within the same day, and across days, weeks, months, and years, revealed consistently high stability in both composition and gene expression within a single day and, importantly, across an approximate six-month period. Among individuals reporting stable diets and no antibiotic exposure, taxonomic and functional profiles remained stable for up to three years. Following colonoscopy preparation, our preliminary study of the microbiome demonstrated strong resilience, returning to its pre-procedure state within one week. Overall, these findings demonstrate that the gut microbiome is generally stable over a six-month time frame, with longer-term changes occurring gradually. These findings support the robustness of stool-based MT profiling for species-level and pathway-resolved functional analysis in longitudinal research and health applications.

## 1. Introduction

Human gut microbiome is a complex community of microorganisms with biochemical activities that are intimately involved in human physiology and health [1–4], like the production and transformation of bioactive molecules [5], modulation of intestinal barrier integrity and mucus biology [6], and calibration of mucosal and systemic immune signaling [7]. Such vital microbial activities are distributed across taxa in various ways; for example, commensal anaerobes that ferment complex carbohydrates and resistant starch can support colonocyte energy metabolism and epithelial homeostasis via butyrate production [8], while mucus-associated taxa participate in mucin turnover and can influence barrier function [9]. Importantly, these microbial activities are dynamic characteristics of the microbiome, raising important questions about how they evolve and adapt over time within an individual.

The extent to which the gut microbiome changes within an individual (intra-donor) across time is still an active area of research [10]. While several studies have investigated gut microbiome stability using sequencing methods, such as 16S and metagenomics, there are limited studies that investigate the transcriptional landscape of the gut microbiome over time [11]. It is important to understand how species-level microbial composition and gene expression profiles are altered within a day, across years, and in response to major luminal perturbations; the findings and physiological insights could inform future studies of the gut microbiome with metatranscriptomics (MT). From a translational perspective, defining expected ranges of variability is also foundational for interpreting microbiome measurements in nutrition, wellness, and disease-adjacent contexts, where a central question is whether an observed change reflects a meaningful biological shift or normal day-to-day fluctuation.

Large metagenomic studies (n>300) have found that intra-individual variability over a range of time scales is lower than inter-individual variability. For example, Mehta *et al*. (n=308) found that the intra-donor gut microbiome composition remained similar, while the metatranscriptome significantly changed after 6 months [11]. Additionally, Chen *et al*. (n=338) found that the gut microbiome and its metabolites remain highly unique to an individual even after four years [12], and Byrd *et al*. (n=413) determined that the gut microbiome and microbial functional potential are stable between 8 and 45 days [13]. These findings support that the gut microbiome may be stable over months and years, though this stability depends on the person’s diet and lifestyle, as well as the analysis methods.

While the previous research has increased our understanding of gut microbiome compositional stability, DNA-based measurements do not investigate alterations in microbial gene expression over time, which MT can provide [14,15]. Alterations in transcriptional activity could occur more quickly than changes in the microbial composition [16]. Therefore, it is important to quantify the stability of the gut microbiome using MT to better understand how stable transcriptional profiles are within an individual over time and in response to perturbation. While microbial composition has been largely shown to remain stable across time using metagenomics, there have been few investigations that directly measure and quantify microbial gene expression profiles within and across days, and none for longer time scales [11].

In addition to the normal microbiome changes, the impact of major luminal perturbations (e.g. colonoscopies) on gut microbiome stability also deserves more thorough investigation. Colonoscopies are becoming increasingly common and have become a mainstay for the diagnosis of colon cancer [17]. Colonoscopies require bowel preparation, which removes the vast majority of the gut microbiome [18]. Prior research showed how the gut microbiome was altered as a consequence of colon lavages [19,20]; with some reports indicating that the gut microbiome recovers within approximately two weeks of a colonoscopy [21–24] and other research indicating long term alterations [25]. Interestingly, there are several preliminary studies implying that the gut dysbiosis caused by lavages could contribute to exacerbating certain conditions, such as Irritable Bowel Syndrome (IBS) [26]. However, all of these studies are in the context of DNA sequencing, and there is a need to better understand how microbial gene expression profiles are altered with colonoscopies.

Characterizing high-resolution microbial composition and gene expression dynamics within the gut microbiome over time – and in response to colonoscopy – can provide valuable insights into transient community shifts and microbial responses to induced perturbation. A key objective of this study is to characterize the stability of the gut microbiome over clinically-relevant time frames. This study investigates the intra-day and long-term (nearly three years) variation in the gut microbiome, as well as its capacity to recover after a colonoscopy using MT.

## 2. Materials and Methods

### 2.1 Ethical conduct of research

This report includes data from two studies. One describes the collection of stool samples pre- and post-colonoscopy and is registered on clinicaltrials.gov (registration number NCT05579444). The other study collected longitudinal data from two participants over almost three years and is also registered in clinicaltrials.gov (registration number NCT06381232). Informed consent was obtained from all subjects involved in the study. The clinical research participants signed informed consent forms, and both studies were approved by an Institutional Review Board (IRB) registered with the United States Health and Human Services. All other analyses described here were performed using customer data and were exempted from an IRB approval and oversight due to the consent by the customers for their data to be used anonymously for research purposes. None of the customers can be identified from these data, as they are aggregated and de-identified, and the sequencing data are not deposited in public repositories.

### 2.2 Study populations

The data presented here were the result of analyses of stool samples from a population of 6,157 individuals. To assess the stability of the gut microbiome over the course of one year, stool samples from 6,150 participants with initial baseline samples and samples collected 1-12 months later were analyzed. The number of participants for each monthly comparison ranged from 81-901. All participants were customers of Viome Life Sciences and received personalized nutritional and supplement recommendations after their baseline samples. Therefore, it is reasonable to expect that the participants may have altered their diet/lifestyle between the baseline and follow-up samples, although diet changes or supplement use were not directly assessed. The nutritional recommendations were not drastic in terms of macronutrient composition, caloric restriction, and/or fasting; they prioritize specific foods for consumption vs. avoidance.

To assess the daily, monthly, and yearly stability of the gut microbiome, two participants collected stool samples over the course of one month, including multiple samples per day, and then another sample almost three years later. During the 2+ years, these participants maintained their diet or lifestyle, did not consume foods with preservatives, and did not take antibiotics, thereby approximating a stable baseline condition without external dietary modulation. This within-subject analysis characterize short-term temporal variability under tightly controlled conditions. The two participants were selected based on their ability to adhere to dense sampling (including multiple samples per day) and maintain highly stable diets and lifestyles over an extended period, thereby minimizing external confounders. While the number of donors is small, the study enables estimation of intra-individual variability across multiple time scales. This approach complements the larger cohort analyses presented here by isolating intrinsic temporal dynamics from inter-individual heterogeneity.

To assess the response of the gut microbiome to a colonoscopy procedure, samples were collected from five participants pre and post colonoscopy. An initial baseline sample was collected prior to colonoscopy and then samples were collected approximately every week after the colonoscopy for four weeks. Diet and lifestyle alterations were not assessed during the study.

### 2.3 Sample collection and metatranscriptomic analysis

All steps were performed using a clinically validated metatranscriptomic method, as previously published [27]. The method includes at-home stool collection, homogenization, and RNA preservation, shipping to the lab, bead beating, nucleic acid extraction, DNase treatment, non-informative RNA depletion, cDNA synthesis, size selection, and limited cycles of PCR for adding dual unique barcodes to each sample. All samples were sequenced using the Illumina NovaSeq 6000 instrument with 2×150 paired-end read chemistry.

### 2.4 Bioinformatic and statistical analyses

Our laboratory maintains a custom reference catalog which includes 32,599 genomes from NCBI RefSeq release 205 ‘complete genome’ category, 4,644 representative human gut genomes of UHGG [28], ribosomal RNA (rRNA) gene sequences, and the human genome GRCh38 [29]. These genomes cover archaea, bacteria, fungi, protozoa, phages, viruses, and the human host. The microbial genomes have 98,527,909 total annotated genes. Our laboratory adopts KEGG Orthology (KO) [30] to annotate microbial gene functions, utilizing the eggNOG-mapper [31] tool for KO assignment.

The clinically validated bioinformatics pipeline maps paired-end reads to the genome catalog using Centrifuge [32] for taxonomy classification (at all taxonomic ranks). Reads mapped to the host genome and rRNA sequences are tracked for monitoring but excluded from further analysis. Reads mapped to microbial genomes are processed with an Expectation-Maximization (EM) algorithm [33] to quantify the relative activity of each microorganism in the sample. Relative activity is equivalent to relative abundance, but it is measured by total readcounts from the transcriptome rather than the genome; it is calculated as the number of transcript reads assigned to a genome, divided by total reads for the sample. Respective taxonomic ranks (strains, species, genera, etc.) can be aggregated from the genomes. For this study, we use species relative activity in the downstream analyses. These genome-mapped reads are further mapped to the gene and open reading frames (ORFs) in order to quantify KO annotated genes or functions.

We define the number of reads mapped to the microbiome as mESD (microbial Effective Sequence Depth) to represent the interpretable portion of the reads in a sample for microbiome. The read counts of identified species and KOs are normalized by mESD in Counts Per Million (CPM) to represent the relative activity. To minimize noise and stochasticity in the data, only species or KOs with minimum 5 reads mapped in each sample were included in downstream analyses.

Inter-donor comparisons were conducted among 408 unique stool samples. The samples were arranged in ascending order based on mESD, and correlation coefficients were calculated between consecutive pairs of samples using interacting species or KOs. This process began with the comparison between the samples with the lowest and second lowest mESD, followed by the third and fourth lowest, and so forth. Correlating samples that attained comparable mESD was done in order to ensure that poor correlation coefficients were not caused by a large discrepancy in the sequencing depth between sample pairs.

All data analysis and statistical tests were performed in Python. ChatGPT-4 Turbo to visualize statistics generated without GenAI.

## 3. Results

### 3.1. Longitudinal stability of the gut microbiome over 12 months

To quantify the longitudinal stability of the gut microbiome, stool was collected from donors at baseline and then again 1-12 months later (Figure 1a). Spearman and Pearson correlation coefficients, along with the Jaccard Index were calculated for microbial composition (species) and functions (KOs) for each donor between baseline and their follow-up collection (Figure 1b). While there is generally high stability of presence (Jaccard), relative activity of species and KOs, further investigation comparing specific longitudinal groups was carried out using these compositional and functional metrics to better understand the influence of time on gut microbiome stability. This involved comparing the correlation coefficients between baseline and 1-month samples (group 1) with the correlation coefficients between baseline and each subsequent month (groups 2-12). For species, there is a decrease in the correlation coefficients between baseline and subsequent months, and these differences become statistically significant after 5 months (for Spearman) to 6 months (for Pearson) (Figure 1c top). A similar trend was observed for KOs, with a decrease in the correlation coefficients between baseline and subsequent months, and these differences become statistically significant after 4 months (for Pearson) to 8 months (for Spearman) (Figure 1c bottom).

**Figure 1.**
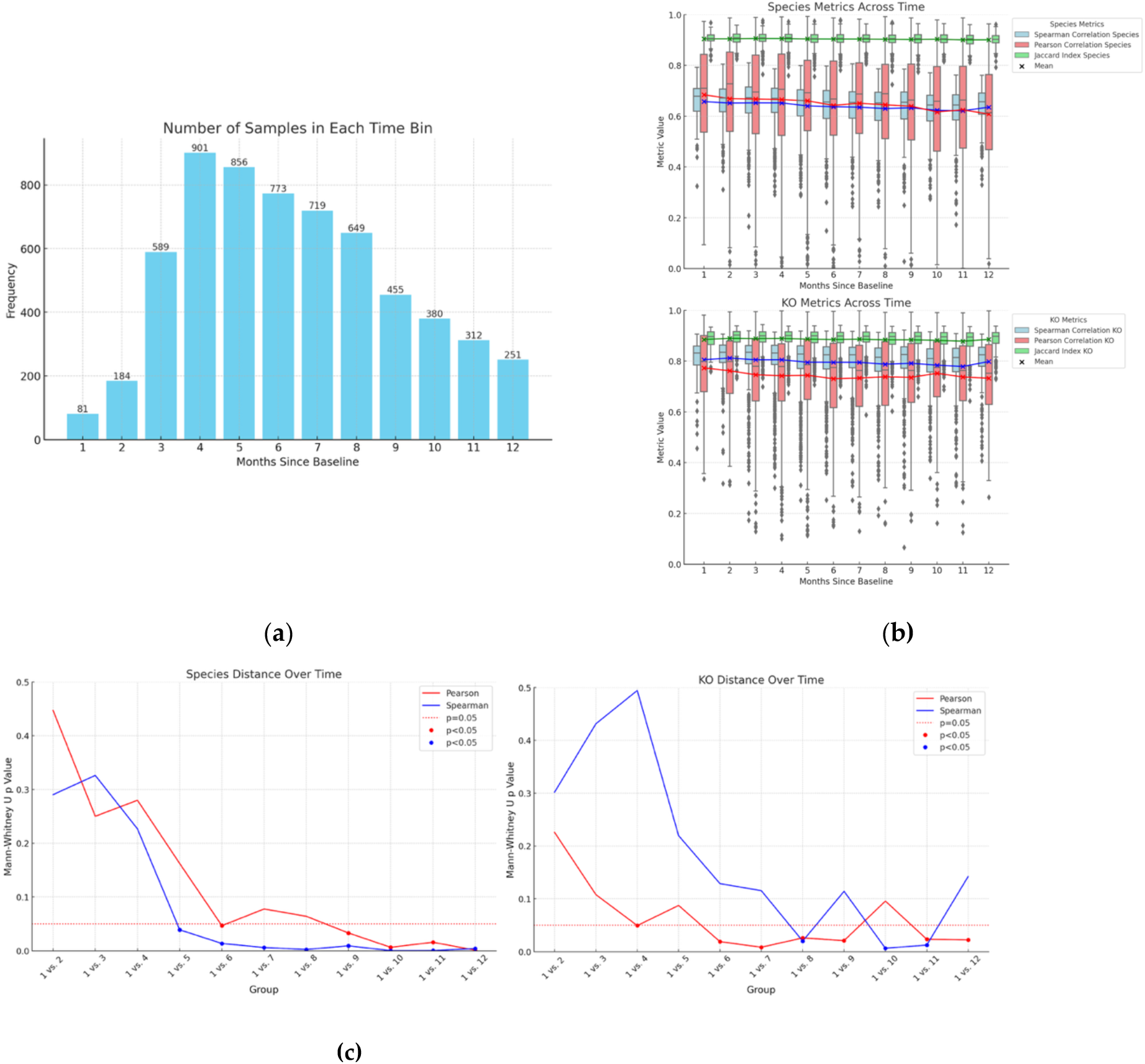
The gut microbiome from 6,150 people shows compositional and functional stability over several months. **(a)** The number of participants for each time period; **(b)** Spearman correlation coefficients, Pearson correlation coefficients, and Jaccard Indices for species and KOs between baseline samples and those collected 1-12 months later; **(c)** p-values from Mann-Whitney U tests, comparing correlation coefficients from month one (vs baseline) to those from months 2-12 (vs baseline). The dotted line represents p=0.05

The Jaccard Indices are generally high for all comparisons: for species, its mean 0.90, and standard deviation 0.03, for KOs, its mean 0.89, and standard deviation 0.04. The sample-specific unique species or KOs had a very small contribution to the overall community composition and functions. For all timepoints, and for both species and KOs, overlapping features had significantly higher relative activities compared to unique features (Supplemental Figure 1, p<0.05).

In summary, although both species and KOs display high stability over a year (Figure 1b), the data demonstrate that the gut microbiome remains highly stable for approximately the first six months, after which correlation coefficients begin a statistically detectable drift (Figure 1c). This drift is gradual and occurs on top of an otherwise stable baseline, indicating that the gut microbiome maintains strong compositional and functional consistency over a six-month time frame. Importantly, participants in this cohort received personalized nutritional recommendations after baseline sampling, which influenced diet and lifestyle. Thus, the gradual drift observed after approximately six months may, at least in part, reflect adaptive responses of the microbiome to these changes rather than a loss of inherent stability under constant conditions.

### 3.2. Exploratory analysis of intra-day, intra-month, and long-term stability

For this analysis, two volunteers (Donors 1 and 2) collected stool samples many times over a period of one month, with two follow-up samples collected at 2 years and 10 months (Donor 1) and then 2 years and 6 months (Donor 2) later. During the month, a total of 9 (Donor 1) and 11 (Donor 2) intra-day samples were collected and compared (Figure 2).

**Figure 2.**
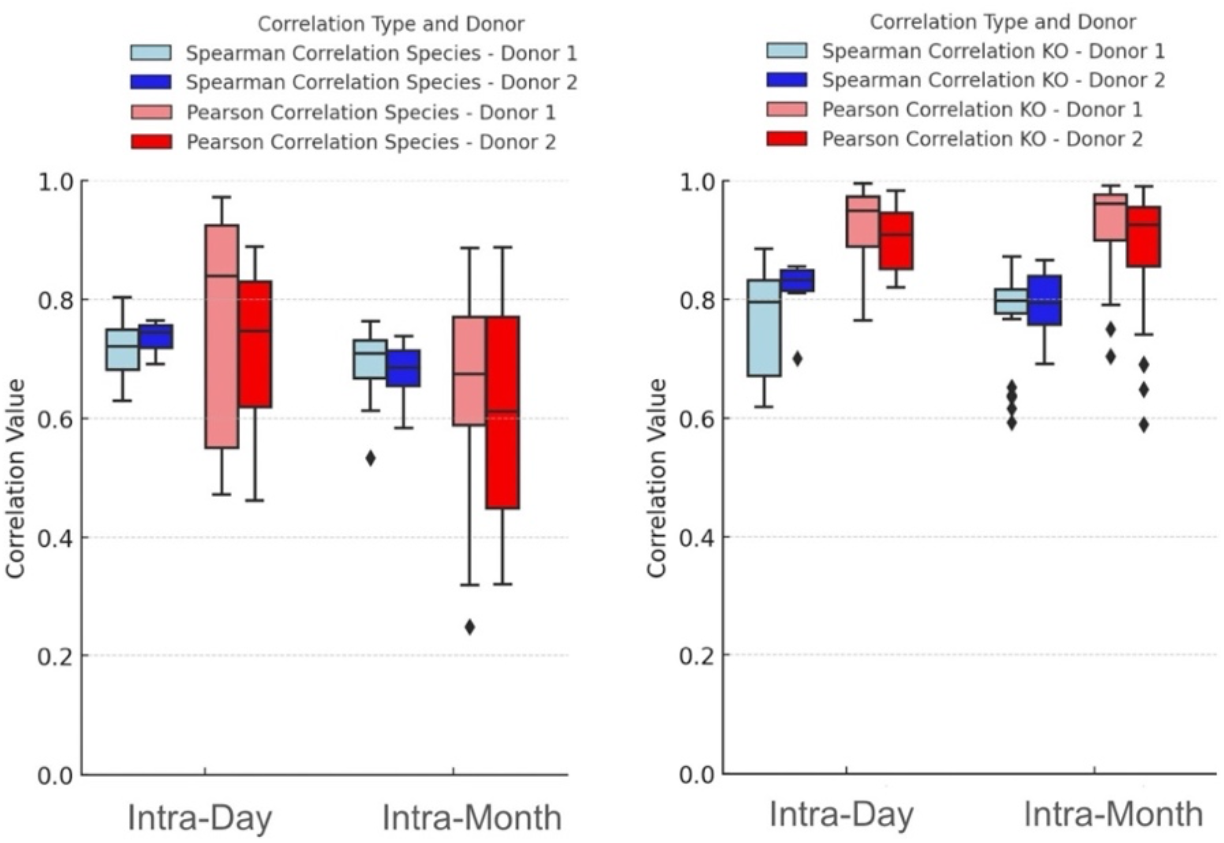
The gut microbiome is stable within a day as well as over the course of a month for taxonomy (species) and microbial functions (KOs). Spearman and Pearson correlation coefficients are shown for microbial species (left panel) and KOs (right panel). Correlation coefficients were significantly higher for the intra-day comparisons compared to the intra-month for species activity (p<0.05).

Spearman and Pearson correlation coefficients were calculated for microbial composition (species, left panel) and functions (KOs, right panel) among all samples collected within a day as well as all those collected within the month. Mann-Whitney U test was used to compare the Spearman and Pearson correlation coefficients between the composite data from the different time points. The Spearman and Pearson correlation coefficients were significantly higher for the intra-day comparisons compared to the intra-month for species activity (p<0.05), indicating that the samples collected on the same day are more highly correlated to one another than those collected over the course of a month for microbial composition. There was no significant difference for Spearman or Pearson correlation coefficients between the intra-day and intra-month groups for KOs (p>0.05), indicating that microbial gene expression is not significantly different among all samples collected within a day as well as among all those collected within the month. However, the stability of functional features (KOs) was high for both intra-day and intra-month, with median values between 0.79 -0.96. The average Jaccard Index for all comparisons for species was 0.763 (standard deviation: 0.038) and 0.746 (standard deviation: 0.069) for KOs. These data show that the gut microbiome composition and functions are stable within a day and over the course of a month, further supporting the broader observation of stability across intermediate time scales up to several months.

Scatter plots correlating the relative activity of species and KOs (Supplemental Figures 2 and 3, for donors 1 and 2, respectively) among samples collected within the same day, one month apart, and almost 3 years apart show reductions in the correlation coefficients and Jaccard Index over time. Although all intra-donor comparisons are significantly correlated (p<0.05) even after almost 3 years, these patterns suggest that there are noticeable changes in the gut microbiome over the course of several years, while also highlighting a surprising degree of stability of an individual’s gut microbiome signature.

**Figure 3.**
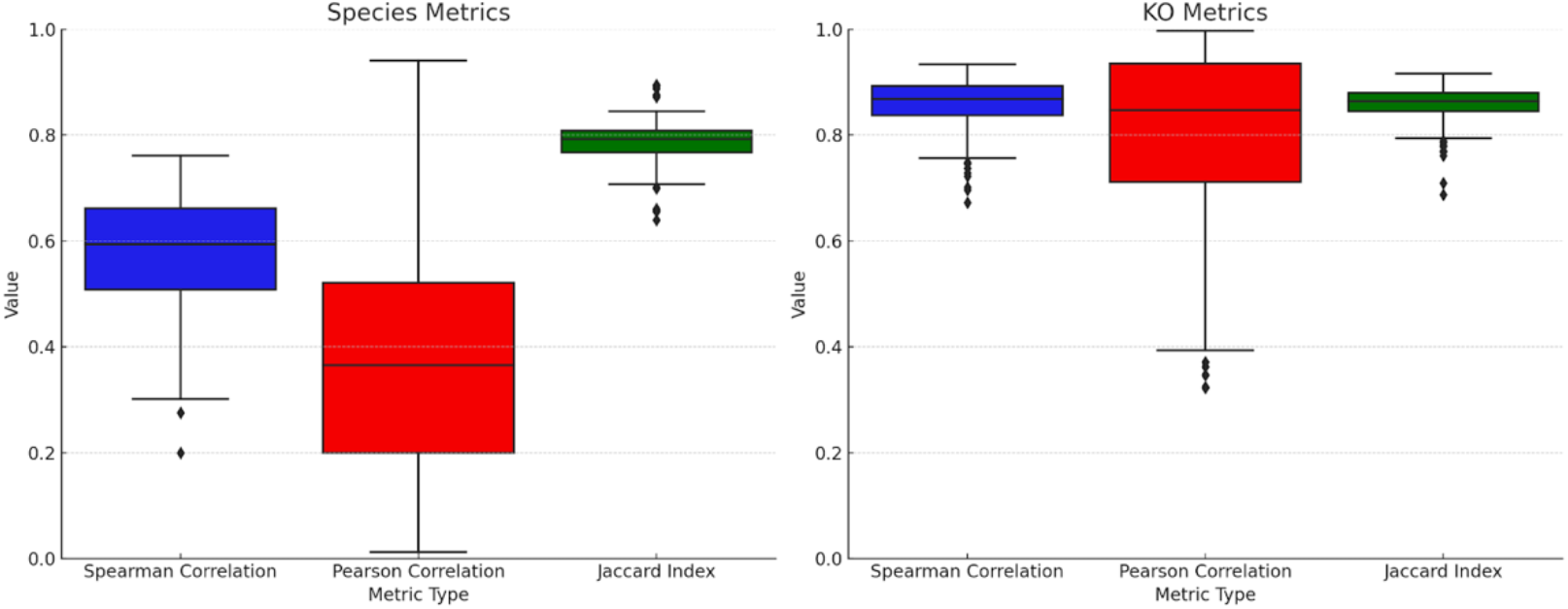
The gut microbiome of different people is dissimilar in terms of species activity but show higher levels of similarity for microbial functions (KOs). Spearman and Pearson correlation coefficients, along with Jaccard Index, are shown for microbial species (left panel) and KOs (right panel) computed from stool analyses from 408 people.

Generally, intra-donor samples that are collected within the same day (intra-day) are more similar to one another than to samples collected on different days (intra-month). However, even over the course of one month, the gut microbiome from the same donor is highly stable in terms of both microbial composition and functions. The taxonomic and functional profile of the gut microbiome is even maintained after almost three years, showcasing high degrees of stability. Interestingly, microbial functions are more highly conserved than microbial taxonomy, and showcase high degrees of similarity across time.

### 3.3. Inter-Donor Comparisons

To better assess the similarity of species and KOs between different individuals, a larger population of inter-donor comparisons were performed (n=408). As seen in Figure 3, species are generally more variable between donors than KOs. These results indicate that microbial composition can be highly donor specific, while microbial functions are more stable across a population

### 3.4. Gut Microbiome Response to Colonoscopy

To determine preliminary stability of the gut microbiome in response to an intestinal perturbation (colonoscopy), stool samples were collected from five participants prior to a colonoscopy (Pre) and then each week after the procedure for four weeks (Post 1-4). Spearman and Pearson correlation coefficients were calculated for microbial composition (species) and functions (KOs), analyzing correlations between pre- and post-samples (Figure 4). The same correlation coefficients were also calculated among all samples from a single donor (intra-donor) as well as among all samples from different donors (inter-donor). All post-colonoscopy samples were significantly correlated with the pre-colonoscopy samples for species and KOs for both Spearman and Pearson correlation coefficients (p<0.05). This indicates the gut microbiome had already recovered by the first week after colonoscopy and was significantly correlated with the pre-colonoscopy community, both in terms of taxonomy and biochemical functions.

**Figure 4.**
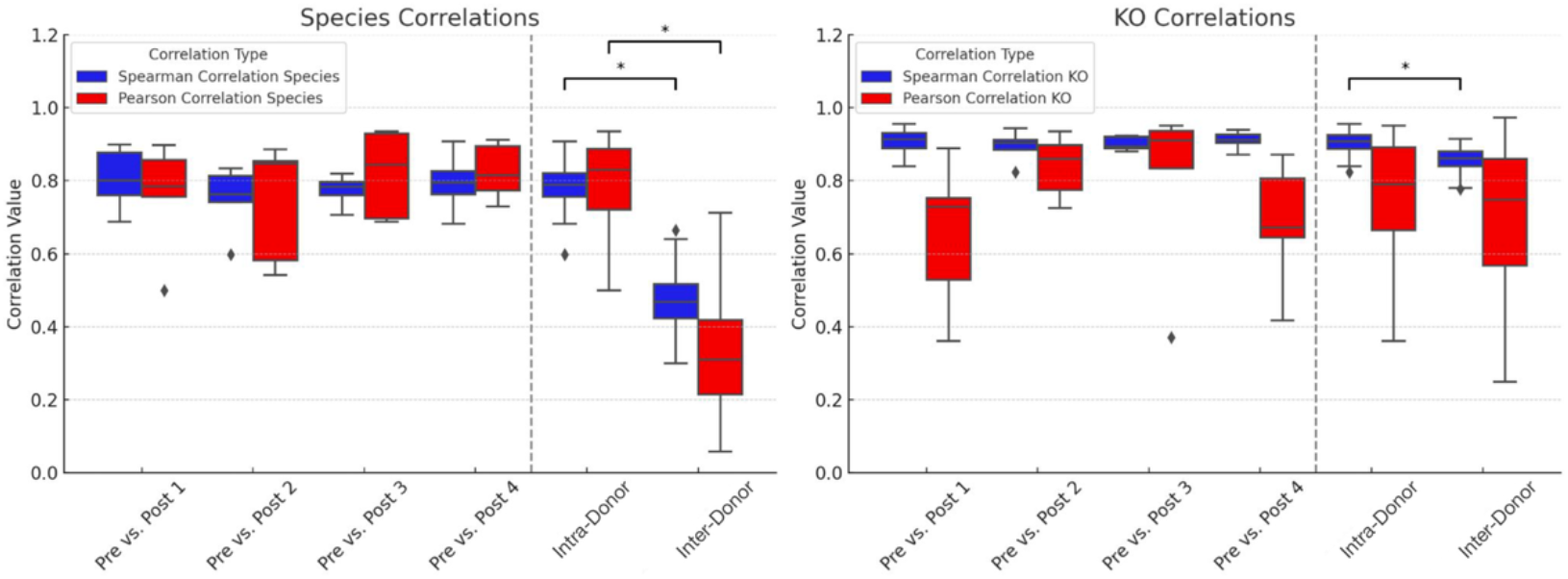
The gut microbiome of different people is dissimilar in terms of species activity but show higher levels of similarity for microbial functions (KOs). Spearman and Pearson correlation coefficients, along with Jaccard Index, are shown for microbial species (left panel) and KOs (right panel) computed from stool analyses from 408 people.

There is a concern that colonoscopies may result in the permanent or long-term loss of intestinal species. To test this hypothesis, Mann-Whitney U tests were used to compare the correlation coefficients between each pair of pre- and post-colonoscopy samples for both species and KOs. There was no significant difference between any of the comparisons (p>0.05). There were significant differences between all intra-donor samples and all inter-donor samples for species (Spearman & Pearson; p<0.05) and for KOs (Spearman; p<0.05), though Pearson did not reach significance for KOs. The average Jaccard Index for all intra-donor comparisons for species was 0.791 (standard deviation: 0.047) and 0.837 (standard deviation: 0.057) for KOs. These results do not provide evidence of substantial loss of species in this small cohort, although larger studies are needed to assess this more definitively. Instead, these preliminary results indicate that the gut microbiome is able to recover from a colonoscopy procedure within approximately one week and that colonoscopies do not dramatically alter the gut microbiome in terms of microbial composition or function.

## 4. Discussion

The gut microbiome, a dynamic and intricate community of microorganisms residing within the gastrointestinal tract, plays a pivotal role in maintaining host health and is increasingly recognized for its involvement in disease pathogenesis [34]. From a physiological perspective, the impact of the microbiome depends not only on the species present but also on their gene expression activities [35]. Thus, characterizing temporal stability at the level of transcription provides important context for how consistently the microbiome may support health-relevant functions over time.

However, the temporal dynamics of the gut microbiome and its response to perturbations, particularly when analyzed by MT, have been understudied. Our study fills this gap by providing valuable insights into the temporal dynamics of the gut microbiome composition and biochemical functions, and their responses to a colonoscopy. Our findings showcase remarkable stability of the gut microbiome within a single day, across several days, and critically, over a period of approximately six months, with longer-term changes occurring more gradually. Our preliminary findings suggest rapid recovery following a colonoscopy, both in terms of taxonomic composition and gene expression. These results underscore the utility of MT to investigate the gut microbiome, offering a powerful lens for understanding how the microbiome adapts and recovers from environmental disruptions.

One of the central findings of our investigation is the long term stability of the gut microbiome, from within a day to almost 3 years (Figures 1 & 2 and Supplemental Figures 2 & 3). Extensive previous metagenomic research has shown that the gut microbiome is relatively stable in terms of microbial composition [11–13], but one transcriptomic study [11] found significant differences after six months, the only time point beyond three days. Our findings provide additional evidence that the stool metatranscriptome changes after six months, on average. However, our data quantify monthly metatranscriptome changes from one to 12 months, providing higher temporal resolution, confirming that it indeed takes 6 months for the stool transcriptome to significantly change, and showing that these changes persist beyond six months. In addition, we show that in two people with a stable diet and lifestyle, and no antibiotic use, the stool metatranscriptome stability can extend beyond two years, which, to our knowledge, is the longest metatranscriptomic study to date. Using MT, our results show that the gut microbiome composition and gene expression profiles are maintained during the day, and remain consistently stable over a period of approximately six months. Physiologically, this pattern is consistent with the idea that the host gut supports a resilient microbial ecosystem capable of maintaining core metabolic activities even as meals and daily routines vary. At the same time, our data show that a baseline gut microbiome begins to drift in terms of both composition and functions after approximately 6 months, highlighting the practical need for retesting when MT is used to track an individual over time.

This drift may reflect, at least in part, exposure to changing dietary inputs rather than an intrinsic temporal instability. Consistent with this interpretation, the two participants who maintained stable diets and lifestyles exhibited sustained microbiome stability for nearly three years, suggesting that, under relatively constant conditions, the gut microbiome may remain stable well beyond six months. The observed slow drift is compatible with gradual changes in diet, seasonality, minor infections, travel, aging, and medication exposures (even when not captured in study metadata), any of which can shift microbial niches and, over time, alter the balance of species and expressed pathways relevant to host physiology. This distinction has important implications for interpreting longitudinal microbiome data. Rather than representing a fixed temporal boundary, the six-month stability window observed here may reflect the timescale over which cumulative dietary and lifestyle modifications begin to measurably reshape microbial community structure and function. In the absence of such changes, our findings suggest that the microbiome can maintain a stable baseline state for substantially longer periods.

A limitation of this study is that the intra-day and intra-month stability analyses are based on only two individuals. Although each participant contributed a large number of longitudinal samples, this small sample size limits the generalizability of these findings. The observed stability may reflect characteristics specific to these individuals, including their controlled diets, stable lifestyles, and absence of antibiotic exposure. Broader populations with more heterogeneous behaviors, health statuses, and environmental exposures may exhibit greater short-term variability. Therefore, these results should be interpreted as a proof-of-concept demonstrating the potential for high temporal stability under controlled conditions, rather than as a definitive estimate of intra-day or intra-month variability across the general population. Another limitation of this study is our inability to compare self-reported phenotypes over time; Viome surveys changed substantially over the years. Therefore we cannot quantify the effects of phenotype changes on the measured microbiome drift presented in this particular study.

Another notable finding is the rapid and robust recovery of the gut microbiome after a colonoscopy (Figure 4). Although these findings are based on a limited sample size (n=5), they align with prior research indicating that colon lavages and other colonoscopy preparation procedures induce temporary disturbances in microbial diversity and composition that recover in a relatively short time span [19–23]. Our analysis adds biochemical functions, which also recover to the pre-colonoscopy state in the first week. This study underscores the resilience of the gut microbiome, which appears well-equipped to quickly restore itself to homeostasis both in terms of microbial composition and functions after major luminal perturbations.

Importantly, our research also shows that while individuals harbor stable microbiomes over time, there is considerable variability between individuals (Figure 3). This duality suggests that while inter-individual microbiomes do exhibit variability, intra-individual microbiomes remain distinctly defined and stable, characterized by a specific set of microbial inhabitants that, on average, maintain a consistent profile over time.

These findings have several implications for both clinical practice and research methodologies. In the clinical realm, our results offer assurance that the transient disruptions induced by colonoscopy preparations are unlikely to have lasting adverse effects on the gut microbiome. Moreover, our study raises the possibility that collecting stool samples from patients shortly after colonoscopy could be feasible, although this observation is based on a small sample size and should be interpreted with caution. Further studies in larger cohorts are needed to determine whether post-colonoscopy sampling can reliably capture representative microbiome states. This approach may streamline sample collection and bolster the assembly of more extensive cohorts for research and diagnostic purposes. In addition, the temporal stability of the gut microbiome, in terms of both species and KO profiles, showcases that stool analyzed with MT is a reliable measure of a stable resident gut microbiome, and not a snapshot of a transient community – an important consideration for designing sampling protocols and interpreting longitudinal monitoring.

Despite the overall stability of the gut microbiome observed in our study, the features directly relevant to human health and disease may be expressed at low levels and, as a result, not adequately reflected in our analyses. Our findings are largely driven by highly expressed housekeeping genes and the activity of highly expressed resident microbial species [36,37].

In conclusion, our MT study demonstrates that the human gut microbiome appears generally stable following major luminal perturbations (e.g. colonoscopy) and over a period of (at least) approximately six months, both in terms of taxonomic composition and gene expression profiles. This stability is observed from intra-day to multi-month time scales, providing a clear window during which the microbiome maintains a consistent baseline state. The gut microbiome composition and functions begin to drift after six months, which indicates that longitudinal testing of the gut microbiome for health applications should be performed after that time period. We also discovered that the gut microbiome appears able to recover from a colonoscopy procedure within 1 week, both in terms of microbial composition and functional capacity.

These results provide a new understanding of the human gut microbiome, which seems to establish a stable composition and gene expression profile that is adapted to the person’s diet. This metatranscriptomic stability does not significantly change after each meal, as others have suggested based on other microbial environments. This gene expression stability may lead to imperfect utilization of all food sources from every meal, but the benefit is that the members of the gut microbiome community do not have to re-establish a new transcriptome after every meal. The gradual drift over longer periods likely reflects the cumulative influence of diet, environment, and host factors on ecological niches in the gut. This combination of short-term stability and longer-term adaptability may be central to the microbiome’s role in host physiology and health.

While this study provides important insights into the temporal stability of the gut microbiome at both taxonomic and functional levels, several avenues for future research could further refine and extend these findings. First, the gradual drift in microbiome composition and function after approximately six months warrants deeper investigation into its underlying drivers. Future studies integrating detailed longitudinal metadata will be essential to disentangle these effects and determine the relative contributions of environmental versus endogenous forces shaping microbiome dynamics. In addition to this, it is clear the intra-day and intra-month stability analyses presented here are based on a limited number of individuals and should be expanded in future studies with dense longitudinal sampling across larger and more diverse cohorts. Finally, an important next step is to establish the clinical relevance of the observed temporal dynamics. Future studies should investigate whether the gradual drift in microbiome composition and function is associated with measurable changes in host physiology or health outcomes. Defining thresholds at which microbiome variation becomes clinically meaningful will be critical for translating metatranscriptomic analyses into actionable diagnostic and monitoring tools. Collectively, these directions will help advance understanding of the mechanisms governing microbiome stability and variability, and they will support the development of more precise and clinically informative microbiome-based applications.

## Supplementary Materials

Figure S1: Mean abundance of shared and unique features, comparing each baseline to monthly sample; Figure S2: Composition and functional profiles for donor 1; Figure S3: Composition and functional profiles for donor 2.

## Author Contributions

Conceptualization, R.T., M.V., G.B.; Data curation, R.T., L.H., N.S.; Formal analysis, R.T., N.S., M.V.; Funding acquisition, M.V., G.B.; Investigation, R.T., L.H., N.S., R.W., M.V., G.B.; Methodology, R.T., L.H., N.S., M.V., G.B.; Project administration, M.V., G.B.; Resources, M.V., G.B.; Software, L.H., N.S.; Supervision, M.V., G.B.; Visualization, R.T., M.V.; Writing – original draft, R.T., M.V.; Writing – review & editing, R.T., L.H., E.P., R.W., M.V., G.B.

## Informed Consent Statement

Informed consent was obtained from all subjects involved in the study.

## Data Availability Statement

The raw data used in this study cannot be shared publicly due to privacy and legal reasons. However, if data is specifically requested, we may be able to share a summary and/or portions of the data. Researchers requiring more data for non-commercial purposes can request via: https://www.viomelifesciences.com/data-access. Viome may provide access to summary statistics through a Data Transfer Agreement that protects the privacy of participants’ data.

## Acknowledgments

During the preparation of this manuscript/study, the author(s) used ChatGPT-4 Turbo to visualize statistics generated without GenAI. The authors have reviewed and edited the output and take full responsibility for the content of this publication.

## Abbreviations

The following abbreviations are used in this manuscript:

MT: Metatranscriptomics
KO: KEGG Orthology

## Supplementary Material

**Figure S1.**
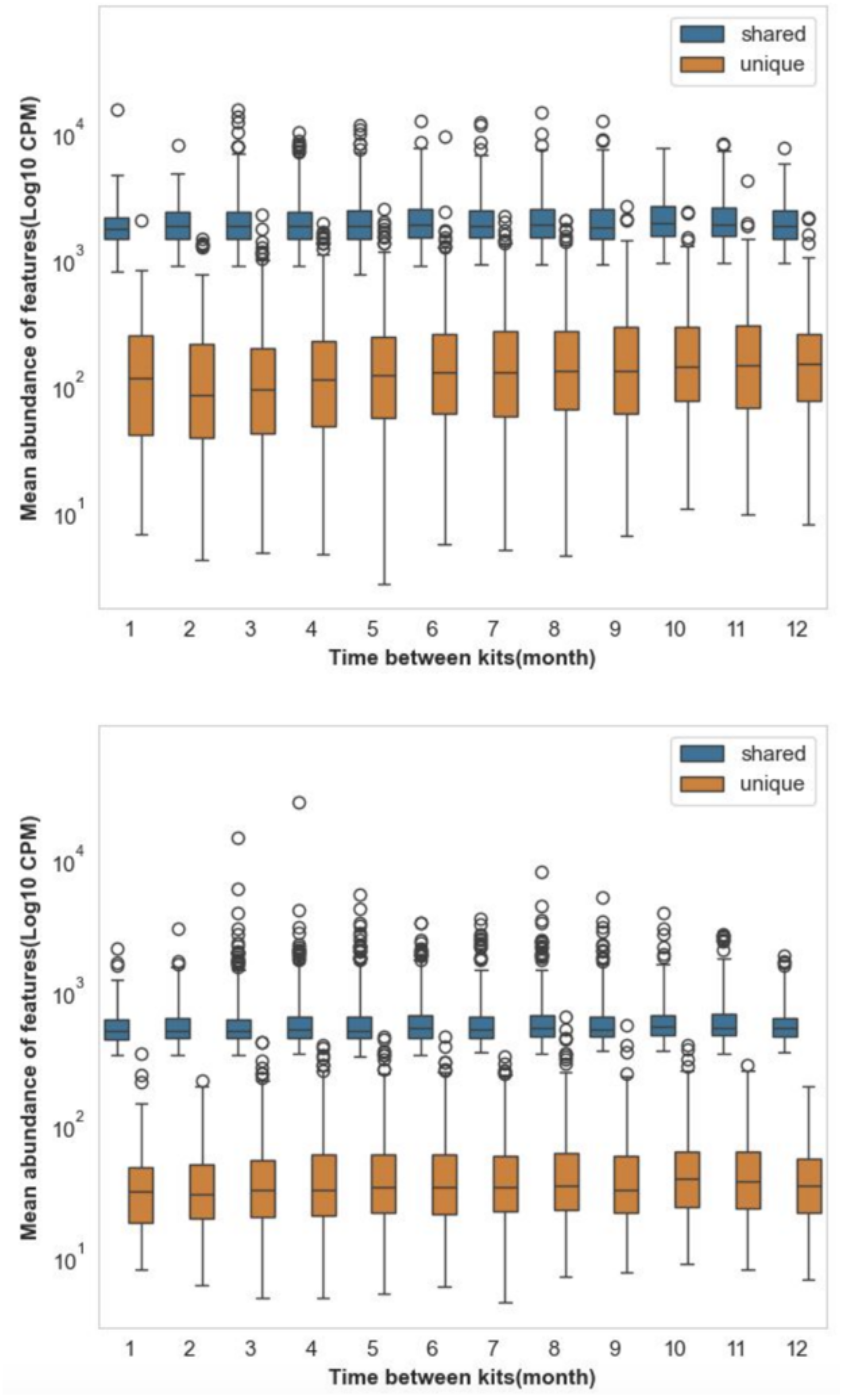
Mean abundance of shared and unique features, comparing each baseline to monthly sample. Shared (blue) and unique (orange) features for species (a) and KOs (b). All months show a significantly (p<0.05) increased abundance of the shared features compared to the unique features.

**Figure S2.**
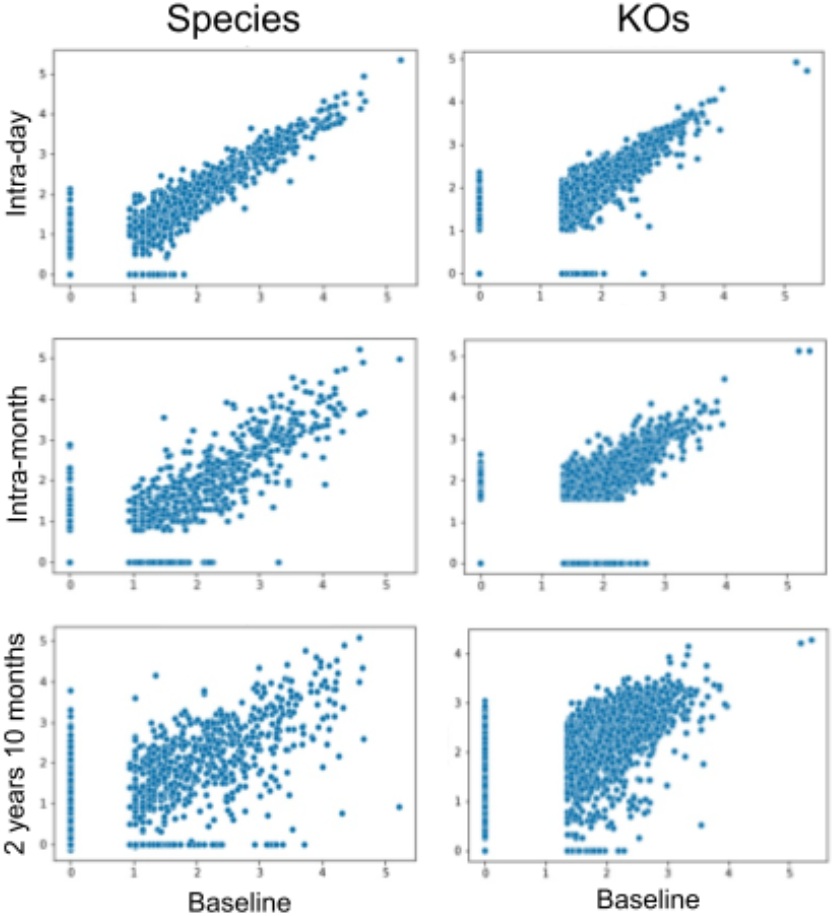
Composition and functional profiles for donor 1. Donor 1 species (left) and KOs (right). The compositional (species) and functional (KO) profiles of the gut microbiome for donor 1 are plotted, normalized to log10(CPM) (Counts Per Million), showing a drift over time. Despite the drift, the gut microbiome retains a significant correlation even after almost three years.

**Figure S3.**
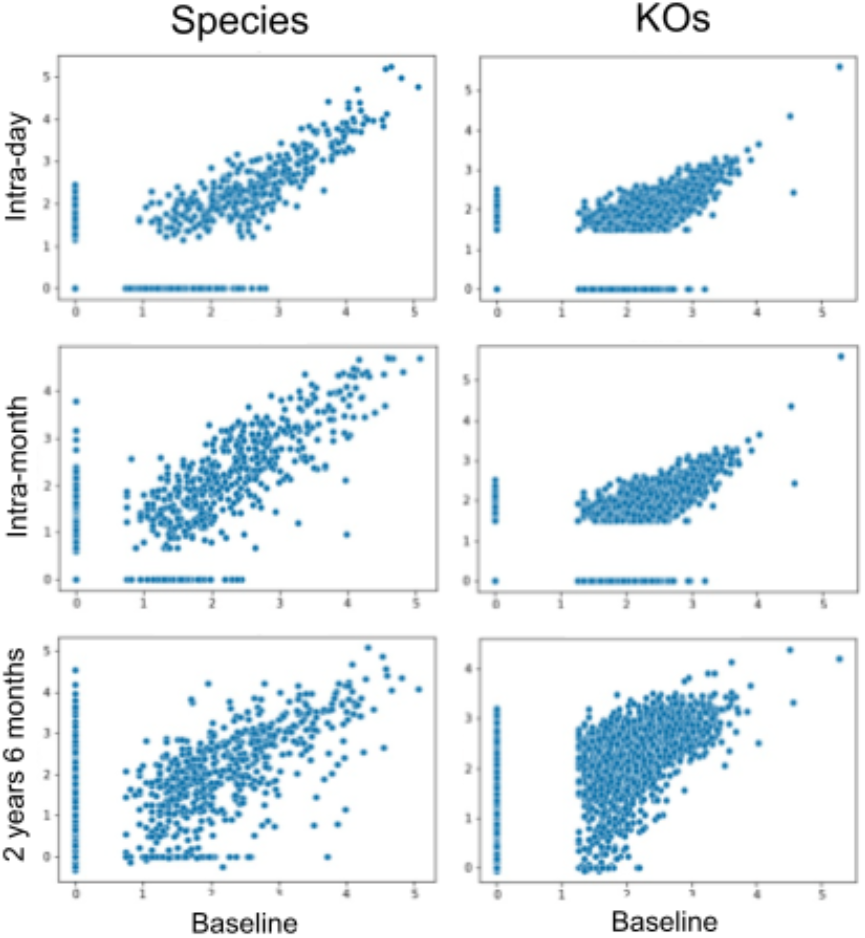
Composition and functional profiles for donor 2. Donor 2 species (left) and KOs (right). The compositional (species) and functional (KO) profiles of the gut microbiome for donor 2 are plotted, normalized to log10(CPM) (Counts Per Million), showing a drift over time. Despite the drift, the gut microbiome retains a significant correlation even after almost three years.

